# Application of class–balancing algorithms to diverse plasma metabolomics datasets using brain tumor as an example

**DOI:** 10.64898/2026.07.02.735756

**Authors:** Adrian Godlewski, Krzysztof Solowiej, Patrycja Mojsak, Joanna Godzien, Julia Zelkowska, Adam Kretowski, Tomasz Lyson, Michal Burdukiewicz, Karol Kaminski, Michal Ciborowski

## Abstract

Class imbalance remains a challenge in metabolomics research, where biological and technical variability can affect statistical inference and machine learning (ML) performance. Class-balancing algorithms address this issue by either increasing minority-class observations or reducing the number of majority-class samples.

This study evaluated the impact of oversampling and undersampling algorithms on targeted and untargeted metabolomics datasets derived from LC-MS and GC-MS analyses of plasma samples from patients with glioblastoma, meningioma, and controls. Synthetic Minority Oversampling Technique (SMOTE) and Random Undersampling (RUS) were applied to balance the datasets, and their effects on data distribution, inter-feature correlations, and machine learning model performance were compared.

RUS preserved the original feature distributions but reduced representativeness by removing the majority-class samples. In contrast, SMOTE introduced synthetic samples that altered covariance structures, increasing the risk of overfitting, particularly in small datasets (n=10). These effects diminished with larger groups (n=30), partially restoring correlations between metabolites. Model performance varied across the class-balancing algorithms. Random Forest classifiers benefited from both balancing methods, with undersampling often yielding higher F1 scores, whereas Support Vector Machine models showed reduced classification performance. These findings highlight the importance of selecting class-balancing strategies based on dataset size, analytical platform, and ML algorithm in metabolomics studies.

## 1. INTRODUCTION

Machine learning (ML) is increasingly used for biomarker discovery and developing strategies to distinguish individuals with diseases from healthy controls [1]. For this purpose, various ML algorithms are trained on omics data, including genomic, proteomic, or metabolomic datasets [2–5]. Omics data are typically characterized by a high-dimensional imbalance [6], which is even more significant in the studies on rare diseases and conditions that are difficult to diagnose early. In these cases, it is challenging to assemble sufficiently large and balanced representative study groups, and comparing a small group with a much larger control population can lead to inaccurate results.

Class-imbalance correction methods are thoroughly documented in the literature and are frequently employed in clinical research [2, 7–11]. Most methods oversample the minority class, while others undersample the majority class or combine both approaches. One of the most widely used oversampling techniques is SMOTE (Synthetic Minority Oversampling Technique), which generates synthetic samples to mitigate overfitting caused by duplicate minority-class samples [10]. SMOTE and its variants have increasingly been applied to balance metabolomic datasets prior to ML model training, potentially improving the classification of minority class samples [8, 11, 12]. Nevertheless, oversampling may introduce noise, distort feature distributions, and increase the risk of false positives, particularly in high-dimensional, low-sample-size datasets [13, 14]. Classification models trained on large numbers of synthetic samples may overfit, compromising model calibration and clinical reliability [13].

Despite numerous methods for addressing class imbalance, clear guidelines for their application in metabolomic studies have yet to be established. The literature reports that applying balancing methods when developing classification models with machine learning algorithms enhances the detection of minority classes [3, 7, 12, 15]. However, some studies suggest that oversampling may introduce noise into the data, adversely affecting statistical inference and increasing the risk of false positives (FP) [13, 14]. Furthermore, classification models trained on large numbers of artificially generated samples may become overfitted and poorly calibrated, leading to misclassification of test subjects. This issue may be particularly relevant in clinical conditions, where incorrect decisions could lead to unnecessary treatment and delays in care for other patients [13]. Despite these potential risks, balance techniques remain valuable tools for preparing data for statistical analysis or modeling. Nevertheless, approaches based on oversampling or undersampling algorithms have not yet been critically evaluated in classification models developed on targeted and untargeted metabolomic data [2, 7, 8, 12, 16–18]. Therefore, this study aimed to evaluate the effectiveness of balancing algorithms and assess their strengths and limitations. We utilized metabolomic data from three complementary techniques with different output data types: targeted analysis by LC-QQQ-MS, untargeted analysis using GC-MS, originating from our earlier published research [19, 20], along with LC-QTOF-MS. The study cohort included patients with glioma and meningioma, as well as a control group, providing a diverse dataset for assessing balancing strategies.

Gliomas are prevalent and aggressive tumors of the central nervous system. These tumors are characterized by considerable heterogeneity, which complicates their classification and early detection, particularly in asymptomatic stages [19]. In contrast, meningiomas generally have a better prognosis, although, like gliomas, they may remain undetected without using imaging techniques [21]. Consequently, assembling a sufficiently large study group to achieve accurate statistical analysis is challenging. Balancing techniques can address this limitation by equalizing the group sizes in datasets, thereby enhancing the ability of ML and statistical tests to detect subtle metabolic differences between study groups.

## 2. MATERIALS AND METHODS

### Design of experiment

To simulate class imbalance, the original datasets were reduced to 10, 20, or 30 observations using Random Undersampling (RUS). The simulated class imbalance was then addressed by applying oversampling techniques, including SMOTE, Adaptive Synthetic Sampling Approach (ADASYN), borderline-SMOTE (BSMOTE), and SMOTE for Numeric and Categorical data (SMOTENC). Additionally, the impact of undersampling, performed using the RUS algorithm to balance group size by reducing the majority group, on modeling was examined. These methods were applied in a loop that iterated across multiple sample sizes. The oversampling process was performed independently for each dataset and each brain tumor group.

To facilitate interpretation of the class distribution in this study, we use the imbalance ratio (IR), defined as the ratio of the number of samples in the majority class to those in the minority class [22]. For both glioma and meningioma, the simulated imbalance conditions corresponded to IR values of 7.10, 3.55, and 2.37 for subsets of 10, 20, and 30 patients, respectively.

To evaluate the effectiveness of oversampling, the Wilcoxon rank sum test (Mann-Whitney U test) was used to compare two independent groups. The p-values for each metabolite were corrected using the Benjamini-Hochberg method. For each group balancing method and sampling condition, Wilcoxon tests were performed in a Monte Carlo simulation with 100 repetitions to reduce the effects of randomness. Each iteration of the algorithm was preceded by limiting the number of observations using the RUS method, ensuring a different, randomly selected set of samples each time. Additionally, the same method was used to test all randomly selected sets without oversampling. Finally, statistics summarizing the Wilcoxon test results were calculated, including the mean, median, standard deviation, and the percentage of cases in which the adjusted p-value of the metabolite fell below the 0.05 significance threshold.

Three different modeling strategies were examined: SMOTE-based oversampling, undersampling (RUS), and unbalanced data (UD). Each strategy follows the same general framework. A subset of glioma or meningioma samples, sized at 10, 20 or 30, was drawn to represent the class with a confirmed disease entity. The three strategies differ in how the training data were assembled. In the oversampling strategy, the minority class was synthetically expanded using SMOTE algorithm with a specified oversampling ratio. The expanded set was then combined with control samples for model training. In the undersampling condition, a random subset of controls was drawn to match the number of case samples, resulting in a balanced but smaller training set. In the UD condition, no resampling was performed, and the original unbalanced subset of case and control samples was used directly. A complementary validation set was constructed by selecting 35 additional case samples and 35 validation controls, resulting in a balanced test set. The trained models were evaluated separately for each condition over 100 iterations. These predictions were compared with the true labels using a confusion matrix to compute standard classification metrics: accuracy, precision, F1 score, Jaccard index, and out-of-sample error (for SVM) or out-of-bag error (for RF). Metrics were recorded for each run, enabling aggregation and comparison across strategies and sample sizes. Figure 1 presents a detailed diagram of the experiment.

**Figure 1.**
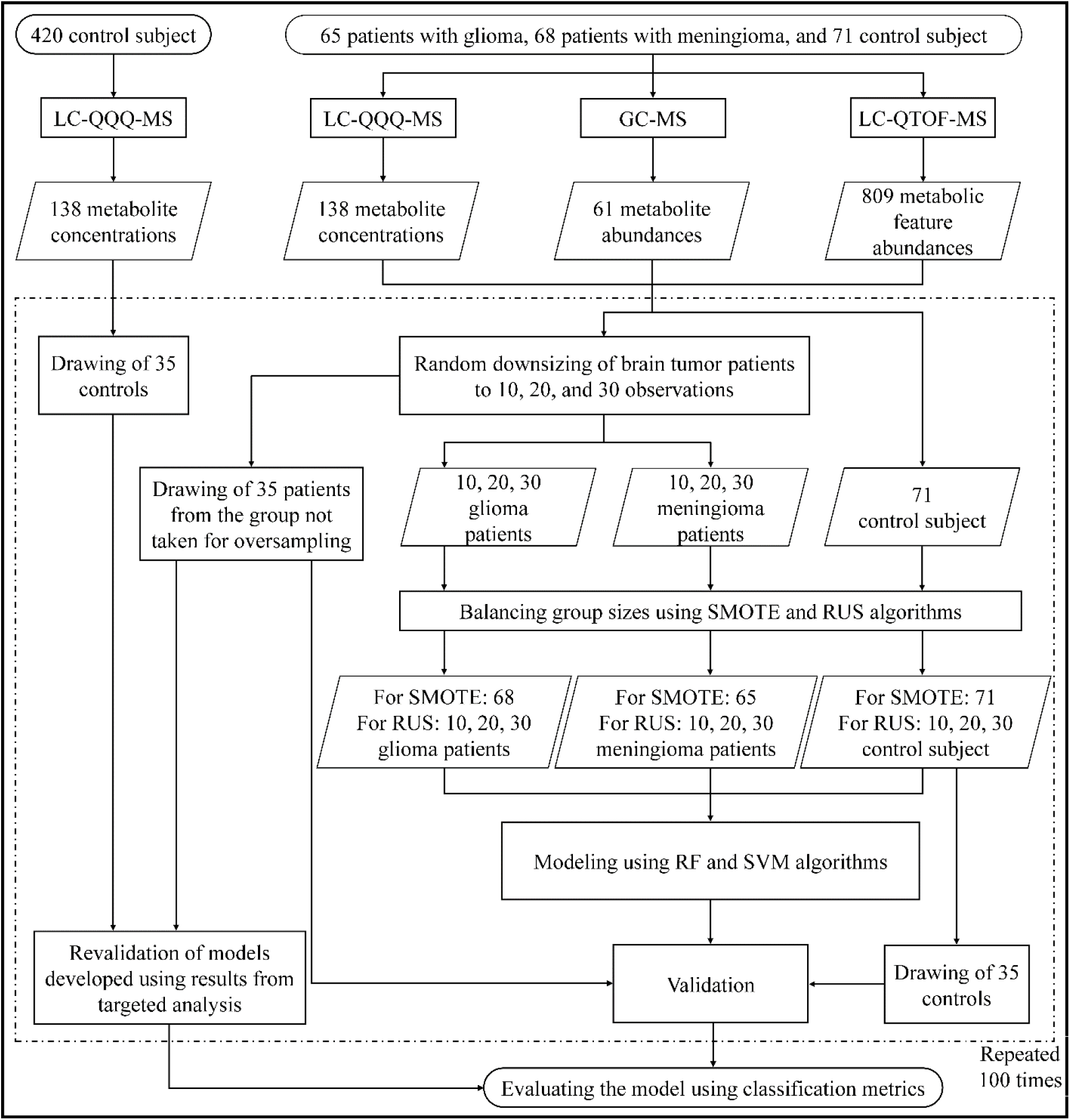
Detailed flowchart of the experiment framework. GC-MS, gas chromatography coupled with mass spectrometry; LC-QQQ-MS, liquid chromatography coupled with tandem mass spectrometry; LC-TOF-MS, liquid chromatography coupled with quadrupole time-of-flight mass spectrometry; RUS, Random Undersampling; SMOTE, Synthetic Minority Over-sampling Technique.

### Patients and groups

The study was conducted using archival samples obtained from the Biobank of the Medical University of Bialystok [23]. Among 202 patients treated at the Neurosurgery Clinic of the University Clinical Hospital in Bialystok between 2016 and 2021, 65 patients with grade 4 glioma and 68 patients with meningioma were included in the study. The tumor malignancy was determined histopathologically according to the fifth edition of the World Health Organization (WHO) classification of tumors of the central nervous system, published in 2021. Based on body mass index (BMI), age, gender, absence of substance abuse, and comorbidities, 491 individuals who underwent routine medical examinations as part of the Bialystok PLUS cohort study were included in the study [24]. In total, the study group consisted of 624 participants, whose characteristics have been previously described [19, 25]. The case and control samples were collected and stored in accordance with biobank standards [26]. The study was approved by the Ethics Committee of the Medical University of Bialystok (No. APK.002.103.2022). Each participant signed an informed consent form before sample collection. The procedures were carried out in accordance with the institutional guidelines of the University Hospital in Bialystok and the Good Clinical Practice guidelines. This study conforms to the principles outlined in the Declaration of Helsinki.

### Targeted metabolomics by LC-QQQ-MS

The targeted analysis of samples has been described previously [20]. In brief, plasma samples for targeted quantification of 188 metabolites were prepared using the AbsoluteIDQ® p180 kit (Biocrates Life Sciences AG, Innsbruck, Austria) according to the manufacturer’s procedure. The metabolites were derivatized with 5% phenyl isothiocyanate in a mixture of ethanol, pyridine, and water (1:1:1, v/v/v). The prepared samples were analyzed using an ultra-high performance liquid chromatograph (1290 Infinity II, Agilent Technologies, USA) coupled with a tandem mass spectrometer (6470 Triple Quad, Agilent Technologies, USA).

### Untargeted metabolomics using GC-MS

The preparation of samples for untargeted analysis has already been described [19]. In brief, plasma samples were simultaneously deproteinized and extracted with 150 µl of cold acetonitrile (1:3, v/v) containing 4-nitrobenzoic acid as an internal standard (ISTD). Metabolite derivatization involved two steps: first, methoxylation with O-methoxyamine hydrochloride in pyridine, followed by a 16-hour incubation at room temperature. Next, samples were silylated by adding N-methyl-N-trimethylsilyl trifluoroacetamide with 1% trimethylchlorosilane. After incubation at 70°C for one hour, 110 µl of heptane containing stearate as a second ISTD was added. The prepared samples were analyzed in random order using a gas chromatograph (7890B series) coupled with a mass spectrometer (7000D series, Agilent Technologies, USA). Metabolites were separated chromatographically on a GC DB-5MS capillary column (30 m × 0.25 mm × 0.25 μm) with helium as the carrier gas.

### Untargeted metabolomics using LC-QTOF-MS

Samples for untargeted metabolomics analysis by LC-QTOF-MS were prepared by adding a cold 1:1 methanol:ethanol mixture containing zomepirac sodium salt as the ISTD to extract metabolites and precipitate plasma proteins. The supernatant was filtered through a 0.22 μm nylon filter into glass chromatography vials [27]. The samples were analyzed in random using liquid chromatograph (1290 Infinity II, Agilent Technologies, USA) coupled with a quadrupole time-of-flight mass spectrometer (6546A, Agilent Technologies, USA) in positive and negative ESI modes.

### Data treatment

Raw spectral data for targeted analysis were processed and quantified using MetIDQ software (Oxygen DB110-3005, Biocrates, Life Science AG, Innsbruck, Austria). Untargeted spectral data from both GC-MS and LC-QTOF-MS were analyzed using Mass Profiler Professional and MassHunter software from Agilent Technologies (USA). Data matrices from the three analytical platforms were normalized and filtered. Only metabolites with a coefficient of variation (CV) in the quality control (QC) samples below 30% for LC-QTOF-MS data and below 35% for GC-MS data were retained. For targeted analysis, metabolites measured in at least 80% of the samples were retained. Missing values in two non-targeted datasets analyses based on LC-QTOF and GC-MS were imputed using the k-nearest neighbors (kNN) algorithm using imputomics application [28]. For targeted analyses, concentrations below the limit of detection (LOD) were replaced with half of the calculated LOD value for each dataset. Before statistical analysis and modeling, the QC samples were excluded. The resulting data matrices were forwarded for further analysis.

### Oversampling and undersampling

The algorithms for balancing group observations were implemented in R (version 4.3.2) using the “themis” package. To generate additional minority class samples, SMOTE, ADASYN, BSMOTE, and SMOTENC were employed. Conversely, the RUS algorithm was used to reduce the number of observations in the majority group. All algorithms aim to balance the sizes of the two groups under study.

### Data analysis

Statistical analyses and graphing were performed using R (version 4.3.2). All comparisons were conducted using the Wilcoxon rank-sum test (Mann-Whitney U test), and p-values for each metabolite were adjusted using the Benjamini-Hochberg method.

To compare the results obtained after oversampling with those from the complete dataset, the terms true positive (TP), false positive (FP), true negative (TN), and false negative (FN) were used as operational categories describing agreement with the complete dataset. The complete dataset was considered the reference set for these comparisons, but not an independently verified biological truth. Under this framework, a TP was defined as a metabolite or metabolic feature identified as statistically significant in both the full and oversampled datasets, whereas a FP was defined as one identified as statistically significant only in the oversampled dataset. Conversely, a TN denoted a metabolite or feature that was not statistically significant in either dataset, and a FN denoted one that was significant in the complete dataset but not in the resampled dataset.

The Jaccard index was calculated as the number of samples classified as TP divided by the total number of observations considered for a specific variant. The Shapiro-Wilk (SW) test was used to assess data distributions for selected metabolites and metabolomic features. The Kolmogorov-Smirnov (KS) test was used to compare the distributions of sets with the balancing algorithms applied to the full set. The use of different analytical techniques enabled correlation analysis for metabolites measured by LC-QQQ-MS and GC-MS, as both platforms provided annotated metabolites. In contrast, LC-QTOF-MS data were not included in this comparison because the detected signals represented anonymous metabolic features that could not be unambiguously assigned to specific metabolites without MS/MS identification. The correlation between metabolites measured by GC-MS and LC-QQQ-MS was assessed using Pearson’s correlation in subgroups where SMOTE was applied. To eliminate dependence among selected samples, each test was performed 100 times, using a different randomly selected subgroup each time.

### Machine learning

The classification models were developed in R using support vector machine (SVM) and random forest (RF) algorithms. Model performance was evaluated through 100 repeated experiments across different training sample sizes. For each iteration, a random subset was used for training, while the remaining samples and validation controls formed the testing set. Models were trained and assessed separately for each sampling method and training size, with predictions made on a fixed, balanced validation set. For SVM, the “caret” package was used with repeated 5-fold cross-validation and a radial basis function (RBF) kernel, which is effective at capturing complex, nonlinear patterns. Each RF model was implemented using the “ranger” package, consisting of 1500 trees, with the number of variables considered at each split set to the square root of the total predictors.

## 3. RESULTS

For clarity and readability, terminology in this section has been simplified. Specifically, LC-QTOF-MS data refer to metabolic features and their abundances, GC-MS data to annotated metabolites measured as abundances, and LC-QQQ-MS data to quantified metabolites reported as concentrations. A statistically significant metabolite or feature (SSM) is defined here as a feature or metabolite whose abundance or concentration differs significantly between the compared groups. Hereafter, unless otherwise stated, the terms “metabolites” are used in a simplified manner, while the platform-specific distinctions described above should be kept in mind.

### Univariate statistics

As previously described [19], GC-MS analysis allowed detection and annotation of 88 metabolites, among which 61 met the quality criteria. LC-QQQ-MS measurement allowed to quantify 188 metabolites, out of which 138 pass the filtration criteria. For untargeted data measured using LC-QTOF-MS, 809 metabolic features remained after quality assurance. These three data matrices were forwarded for statistical analysis. In the full dataset comparing gliomas and controls, the U test revealed 292 SSM for LC-QTOF-MS, 38 SSM for GC-MS, and 70 SSM for LC-QQQ-MS data. For the comparison between meningiomas and controls, 174 SSM for LC-QTOF-MS, 41 SSM for GC-MS and 19 SSM for LC-QQQ-MS were selected.

Firstly, a U test was performed to compare data processed with four oversampling algorithms (SMOTE, BSMOTE, ADASYN, and SMOTENC). Based on the average number of SSM across 100 iterations, it was concluded that SMOTE, ADASYN, and SMOTENC give similar results (see Supplementary Figure S1). Conversely, the BSMOTE algorithm failed when applied to small groups. Since the SMOTE algorithm is widely used in metabolomics and other omics studies, we chose it as the representative algorithm for this group of balancing methods [3, 8, 11, 17, 29]. The U test, along with the Benjamin-Hochberg correction, was used to evaluate various approaches to addressing the imbalance between the study groups. The results of the comparison between SMOTE, RUS, UD, and the full data set are presented in Figure 2. The p-values from the U tests were further analyzed, enabling the selection of metabolites incorrectly labeled as statistically significant compared with the full data set. The calculated median values for SSM, FPs, and overlapping metabolites across datasets for 100 iterations are presented in Table 1.

**Figure 2.**
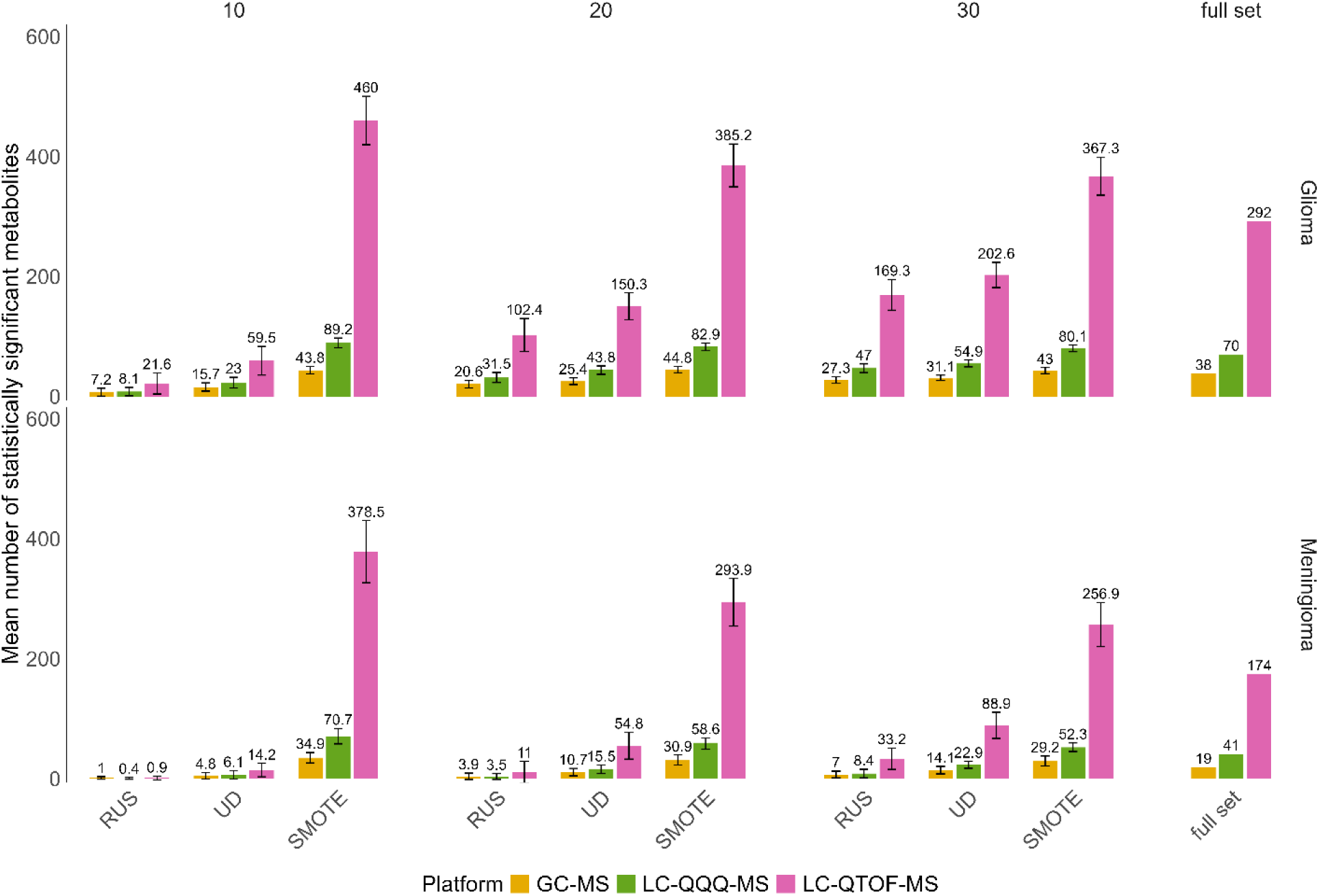
The average number of SSM for GC-MS, LC-QQQ-MS, and LC-QTOF-MS, across 100 iterations, considering subgroup division and the application of balancing algorithms. GC-MS, gas chromatography coupled with mass spectrometry; LC-QQQ-MS, liquid chromatography coupled with tandem mass spectrometry; LC-TOF-MS, liquid chromatography coupled with quadrupole time-of-flight mass spectrometry; RUS, Random Undersampling; SMOTE, Synthetic Minority Over-sampling Technique; UD, unbalanced data.

**Table 1.** Comparison of three analytical platforms based on median values of true positive and false positive metabolic features and their F1 scores.

| N | Method | LC-QQQ-MS |  |  | GC-MS |  |  | LC-QTOF-MS |  |  |
| --- | --- | --- | --- | --- | --- | --- | --- | --- | --- | --- |
|  |  | TP | FP | F1 score | TP | FP | F1 score | TP | FP | F1 score |
| Glioma |  |  |  |  |  |  |  |  |  |  |
| 10 | UD | 18 | 0 | 0.41 | 12 | 0 | 0.48 | 36 | 0 | 0.22 |
| 20 |  | 39 | 0 | 0.72 | 23 | 0 | 0.75 | 130 | 0 | 0.62 |
| 30 |  | 54 | 0 | 0.87 | 30 | 0 | 0.88 | 193 | 0 | 0.80 |
| 10 | SMOTE | 69 | 16 | 0.89 | 38 | 10 | 0.88 | 292 | 117 | 0.83 |
| 20 |  | 69 | 12 | 0.91 | 38 | 8 | 0.90 | 291 | 52 | 0.92 |
| 30 |  | 69 | 11 | 0.92 | 38 | 5 | 0.94 | 292 | 56 | 0.91 |
| 10 | RUS | 3 | 0 | 0.08 | 2 | 0 | 0.10 | 4 | 0 | 0.03 |
| 20 |  | 25 | 0 | 0.53 | 20 | 0 | 0.69 | 78 | 0 | 0.42 |
| 30 |  | 46 | 0 | 0.79 | 26 | 0 | 0.81 | 144 | 0 | 0.66 |
| Meningioma |  |  |  |  |  |  |  |  |  |  |
| 10 | UD | 2 | 0 | 0.09 | 2 | 0 | 0.19 | 3 | 0 | 0.03 |
| 20 |  | 8 | 0 | 0.33 | 8 | 0 | 0.59 | 32 | 0 | 0.31 |
| 30 |  | 20 | 0 | 0.66 | 10 | 0 | 0.69 | 59 | 0 | 0.51 |
| 10 | SMOTE | 39 | 18 | 0.80 | 19 | 15 | 0.72 | 171 | 97 | 0.77 |
| 20 |  | 40 | 11 | 0.87 | 19 | 9 | 0.81 | 172 | 50 | 0.87 |
| 30 |  | 41 | 8 | 0.91 | 19 | 7 | 0.84 | 170 | 45 | 0.87 |
| 10 | RUS | 0 | 0 | 0.00 | 0 | 0 | 0.00 | 0 | 0 | 0.00 |
| 20 |  | 0 | 0 | 0.00 | 1 | 0 | 0.10 | 2 | 0 | 0.02 |
| 30 |  | 3 | 0 | 0.14 | 4 | 0 | 0.35 | 17 | 0 | 0.18 |

FP, false positive; GC-MS, gas chromatography coupled with mass spectrometry; LC-QQQ-MS, liquid chromatography coupled with tandem mass spectrometry; LC-QTOF-MS, liquid chromatography coupled with quadrupole time-of-flight mass spectrometry; RUS, Random Undersampling; SMOTE, Synthetic Minority Over-sampling Technique; TP, true positive; UD, unbalanced data.

When comparing results for groups with different imbalance ratios (7.10, 3.55, and 2.37), the ratio of the mean number of statistically significant metabolites in the UD to the total number of SSM in the full set decreases as the number of measured metabolites increases. In the UD of 10 glioma patients obtained using LC-QTOF-MS, only 20.4% of metabolic features are significant compared with the full set, whereas in GC-MS data, this figure is 41.3%. Importantly, in the UD of meningiomas, a considerably lower percentage of overlapping SSM with the full set was obtained compared with UD gliomas, regardless of the analytical platform (see Table 1).

However, processing the data with the SMOTE algorithm led to a substantial increase in the number of metabolites across all tested groups and platforms. Notably, oversampling enabled the identification of nearly all significant changes observed in the full set. However, this approach led to a significant increase in FPs, as evidenced by a mean of 117 FPs in the LC-QTOF-MS dataset comprising 10 glioma samples (Table 1). The number of FPs decreased as the number of samples subjected to SMOTE balancing increased. Applying RUS to balance group sizes eliminated FPs, but the average number of metabolites across all data sets was significantly lower than with UD (see Table 1). In glioma sets, the decrease in significant changes was -63.7%, -31.9%, and -16.4% for group sizes of 10, 20, and 30, respectively. For the meningioma sets, the reduction was -93.6%, -79.9%, and -62.7%, respectively (see Figure 2).

### Data distribution analysis

KS test was performed to assess how the use of class-balancing algorithms affects data distributions. The median p-values are presented in Figure 3A for SMOTE and in Figure 3B for RUS. In highly unbalanced subgroups, applying SMOTE distorted the dataset’s similarity relative to the full dataset. This effect was independent of the analytical platform used. Interestingly, for GC-MS, the glioma subgroup had a significantly lower p-value than the meningioma subgroup. To better illustrate the distribution changes introduced by SMOTE, violin plots were prepared for selected metabolites representing different distribution patterns (see Figure 4). The slight variation in leucine abundances obtained by GC-MS in the glioma group precludes reproducing the data distribution from the full set. An SW test was performed, which revealed a change in the distribution of leucine data. A subset of 10 glioma patients, balanced by SMOTE, exhibited a non-normal distribution (Supplementary Table S1), whereas in the full set, this metabolite showed a normal distribution (Supplementary Table S2). For other metabolites, p-values were close to the statistical significance threshold, but no distributional change was detected (Supplementary Table S1 and S2). In contrast, using RUS yielded distributions more similar to those of the full dataset, even with a small study group (Figure 3B). Violin plots for selected metabolites and metabolic features, showing groups balanced using the RUS algorithm, are presented in Supplementary Figure S2.

**Figure 3.**
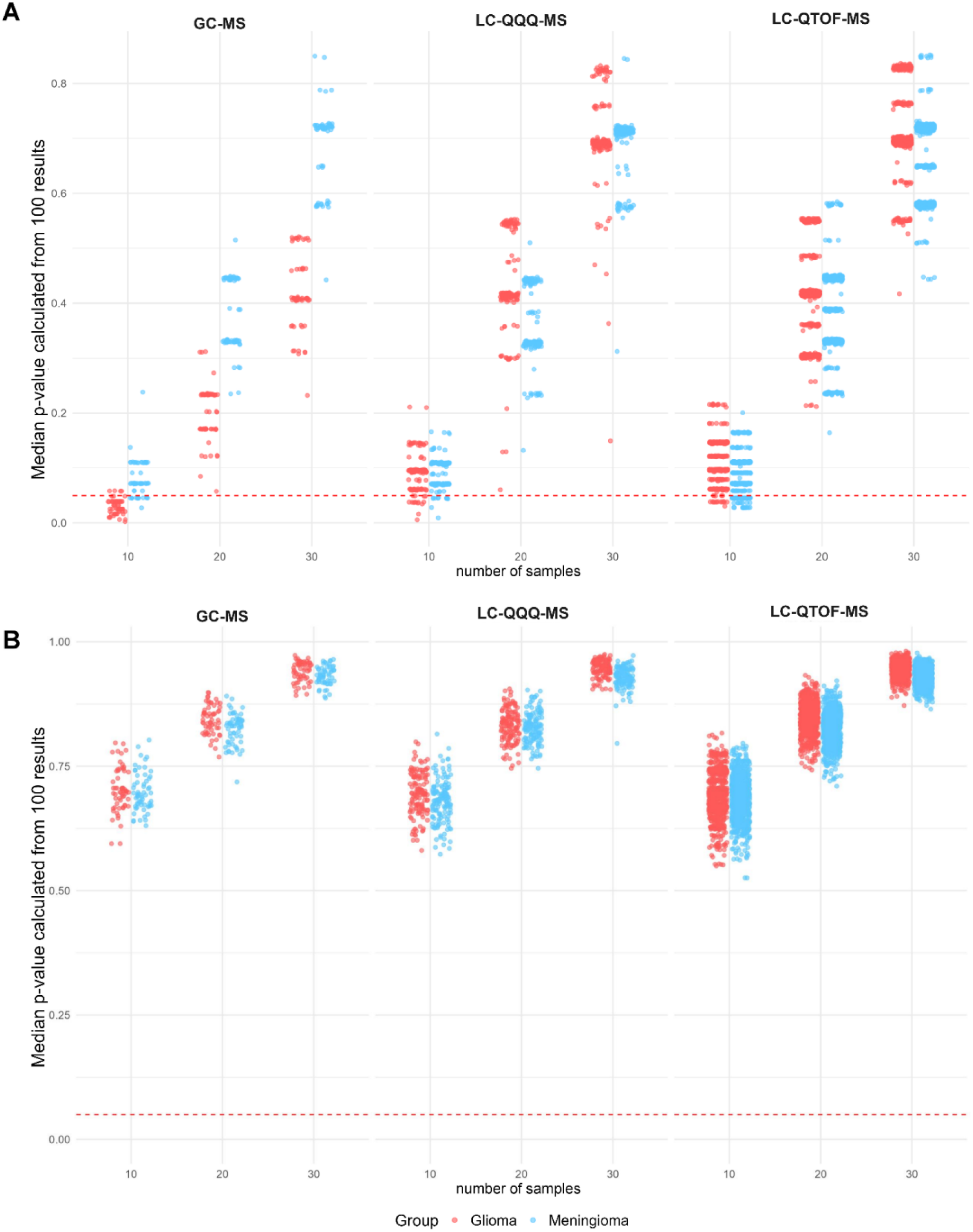
Median p-values from the Kolmogorov-Smirnov test for glioma and meningioma subsets balanced using (A) SMOTE and (B) RUS. GC-MS, gas chromatography coupled with mass spectrometry; LC-QQQ-MS, liquid chromatography coupled with tandem mass spectrometry; LC-QTOF-MS, liquid chromatography coupled with quadrupole time-of-flight mass spectrometry.

**Figure 4.**
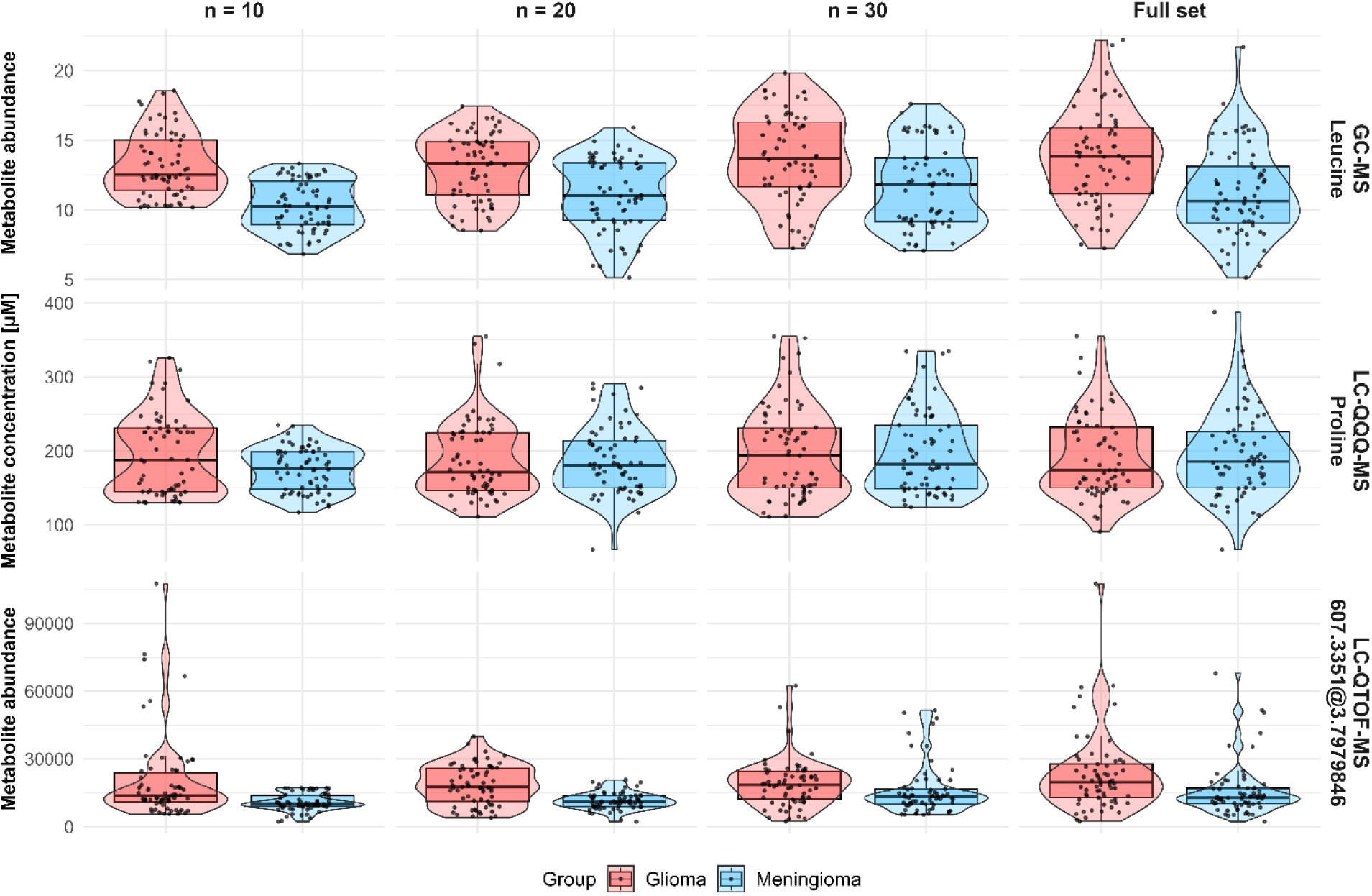
Violin plots representing the abundance of leucine measured by GC-MS, the concentration of proline measured by LC-QQQ-MS, and the abundance of the metabolic feature with a neutral mass of 607.3351 Da measured by LC-QTOF-MS after balancing the groups using SMOTE. GC-MS, gas chromatography coupled with mass spectrometry; LC-QQQ-MS, liquid chromatography coupled with tandem mass spectrometry; LC-QTOF-MS, liquid chromatography coupled with quadrupole time-of-flight mass spectrometry.

### Metabolite correlations

Correlation analysis across 10, 20, 30 subgroups and the full set showed that using SMOTE to balance group sizes distorts the correlation results (see Figure 5). The p-value indicated that, for 10 samples, only 3 metabolites exhibit a statistically significant correlation, whereas in the full set, 11 metabolites show a strong correlation. However, when applying SMOTE to a subgroup of 30 patients, the correlation strength was similar to that in the full set.

**Figure 5.**
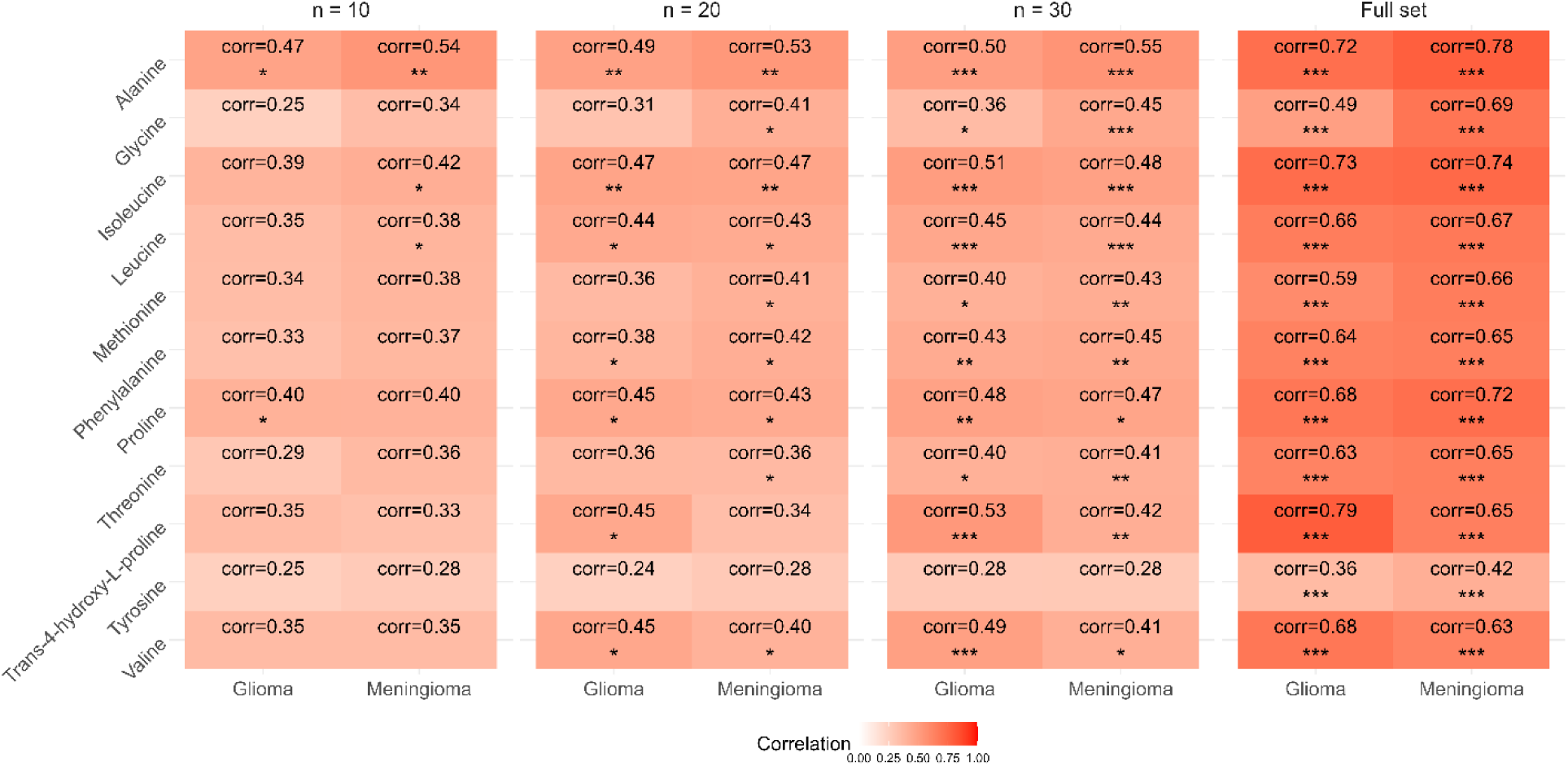
Heatmap illustrating Pearson’s correlation of the same metabolites measured by GC-MS and LC-QQQ-MS for the full set and subsets of 10, 20, and 30 balanced using SMOTE. Significance levels are indicated as follows: * p-value <0.05; ** p-value < 0.01; *** p-value < 0.001.

### Machine learning modeling

Data sets balanced using SMOTE, RUS, and UD were used to develop RF and SVM models, and their performance is summarized in Table 2. For RF-based models, applying SMOTE increased the average F1 score by 12.3% (8.7–15.2%) for gliomas and 35.2% (14.3–61.0%) for meningioma across three platforms, compared with models built on unbalanced data. However, using RUS to balance the groups increased the average F1 score by 19.6% (15.7–24.0%) for glioma and 87.0% (40.6–148.4%) for meningioma. Conversely, for SVM models trained with SMOTE, a decline in the F1 score was observed for both glioma (average percentage change: -4.6%) and meningioma (average percentage change: -5.5%). Notably, the use of RUS increased the F1 score for subsets of 10 samples, whereas groups of 20 and 30 patients showed either a decrease or only a slight increase in both F1 score and accuracy. Untargeted metabolomic data obtained using LC-QTOF-MS, combined with RUS, enabled the development of the best models. Models trained on unbalanced data had lower average F1 scores than models supported by RUS, as evidenced by average percentage changes of 42.9% for RF and 17.3% for SVM. However, balancing groups with RUS resulted in an average decrease in precision of about 14.5% for both the RF-based and SVM-based models.

**Table 2.** The mean values (n = 100) of the classification metrics computed for models trained with RF and SVM.

| N | Parameter | Targeted analysis - LC-QQQ-MS |  |  | Untargeted analysis - GC-MS |  |  | Untargeted analysis - LC-QTOF |  |  |
| --- | --- | --- | --- | --- | --- | --- | --- | --- | --- | --- |
|  |  | UD | SMOTE | RUS | UD | SMOTE | RUS | UD | SMOTE | RUS |
| Random Forest |  |  |  |  |  |  |  |  |  |  |
| Glioma |  |  |  |  |  |  |  |  |  |  |
| 10 | Accuracy | 0.74 | 0.81 | 0.86 | 0.69 | 0.78 | 0.84 | 0.67 | 0.79 | 0.90 |
|  | F1 score | 0.64 | 0.76 | 0.86 | 0.54 | 0.71 | 0.83 | 0.51 | 0.74 | 0.90 |
|  | Jaccard index | 0.48 | 0.62 | 0.75 | 0.38 | 0.56 | 0.71 | 0.35 | 0.59 | 0.81 |
| 20 | Accuracy | 0.84 | 0.88 | 0.90 | 0.81 | 0.87 | 0.87 | 0.85 | 0.91 | 0.93 |
|  | F1 score | 0.80 | 0.86 | 0.89 | 0.76 | 0.84 | 0.86 | 0.82 | 0.90 | 0.93 |
|  | Jaccard index | 0.68 | 0.75 | 0.81 | 0.61 | 0.73 | 0.76 | 0.70 | 0.82 | 0.87 |
| 30 | Accuracy | 0.88 | 0.90 | 0.92 | 0.86 | 0.88 | 0.87 | 0.91 | 0.94 | 0.94 |
|  | F1 score | 0.86 | 0.88 | 0.91 | 0.84 | 0.87 | 0.87 | 0.90 | 0.94 | 0.94 |
|  | Jaccard index | 0.75 | 0.80 | 0.84 | 0.73 | 0.77 | 0.77 | 0.83 | 0.88 | 0.89 |
| Meningioma |  |  |  |  |  |  |  |  |  |  |
| 10 | Accuracy | 0.54 | 0.59 | 0.69 | 0.58 | 0.61 | 0.67 | 0.52 | 0.55 | 0.74 |
|  | F1 score | 0.15 | 0.29 | 0.67 | 0.28 | 0.36 | 0.64 | 0.08 | 0.19 | 0.72 |
|  | Jaccard index | 0.07 | 0.18 | 0.51 | 0.17 | 0.22 | 0.48 | 0.03 | 0.10 | 0.57 |
| 20 | Accuracy | 0.63 | 0.71 | 0.76 | 0.68 | 0.72 | 0.71 | 0.59 | 0.70 | 0.80 |
|  | F1 score | 0.41 | 0.59 | 0.75 | 0.52 | 0.62 | 0.69 | 0.31 | 0.57 | 0.79 |
|  | Jaccard index | 0.26 | 0.42 | 0.60 | 0.35 | 0.45 | 0.53 | 0.18 | 0.41 | 0.66 |
| 30 | Accuracy | 0.71 | 0.79 | 0.81 | 0.74 | 0.76 | 0.73 | 0.69 | 0.81 | 0.84 |
|  | F1 score | 0.59 | 0.72 | 0.81 | 0.65 | 0.69 | 0.70 | 0.56 | 0.76 | 0.84 |
|  | Jaccard index | 0.42 | 0.57 | 0.68 | 0.48 | 0.53 | 0.55 | 0.39 | 0.62 | 0.72 |
| Support Vector Machine |  |  |  |  |  |  |  |  |  |  |
| Glioma |  |  |  |  |  |  |  |  |  |  |
| 10 | Accuracy | 0.82 | 0.80 | 0.88 | 0.74 | 0.70 | 0.83 | 0.75 | 0.67 | 0.87 |
|  | F1 score | 0.78 | 0.73 | 0.88 | 0.65 | 0.57 | 0.83 | 0.67 | 0.50 | 0.86 |
|  | Jaccard index | 0.65 | 0.59 | 0.79 | 0.48 | 0.41 | 0.71 | 0.51 | 0.34 | 0.76 |
| 20 | Accuracy | 0.91 | 0.90 | 0.92 | 0.87 | 0.87 | 0.86 | 0.90 | 0.87 | 0.92 |
|  | F1 score | 0.90 | 0.88 | 0.92 | 0.85 | 0.85 | 0.86 | 0.89 | 0.85 | 0.91 |
|  | Jaccard index | 0.82 | 0.79 | 0.85 | 0.74 | 0.74 | 0.76 | 0.80 | 0.74 | 0.84 |
| 30 | Accuracy | 0.93 | 0.93 | 0.94 | 0.91 | 0.91 | 0.89 | 0.94 | 0.93 | 0.93 |
|  | F1 score | 0.93 | 0.93 | 0.94 | 0.90 | 0.90 | 0.88 | 0.94 | 0.93 | 0.93 |
|  | Jaccard index | 0.87 | 0.86 | 0.89 | 0.82 | 0.83 | 0.79 | 0.88 | 0.87 | 0.87 |
| Meningioma |  |  |  |  |  |  |  |  |  |  |
| 10 | Accuracy | 0.58 | 0.60 | 0.71 | 0.57 | 0.57 | 0.69 | 0.52 | 0.52 | 0.72 |
|  | F1 score | 0.31 | 0.33 | 0.69 | 0.32 | 0.25 | 0.66 | 0.17 | 0.11 | 0.71 |
|  | Jaccard index | 0.15 | 0.20 | 0.53 | 0.14 | 0.15 | 0.50 | 0.04 | 0.04 | 0.56 |
| 20 | Accuracy | 0.77 | 0.76 | 0.79 | 0.74 | 0.72 | 0.74 | 0.74 | 0.69 | 0.82 |
|  | F1 score | 0.70 | 0.69 | 0.79 | 0.65 | 0.61 | 0.73 | 0.64 | 0.55 | 0.82 |
|  | Jaccard index | 0.54 | 0.53 | 0.66 | 0.49 | 0.45 | 0.58 | 0.48 | 0.38 | 0.69 |
| 30 | Accuracy | 0.86 | 0.85 | 0.84 | 0.80 | 0.82 | 0.78 | 0.85 | 0.82 | 0.86 |
|  | F1 score | 0.84 | 0.83 | 0.84 | 0.74 | 0.78 | 0.77 | 0.82 | 0.78 | 0.86 |
|  | Jaccard index | 0.73 | 0.71 | 0.72 | 0.59 | 0.64 | 0.63 | 0.69 | 0.64 | 0.76 |

GC-MS, gas chromatography coupled with mass spectrometry; LC-QQQ-MS, liquid chromatography coupled with tandem mass spectrometry; LC-QTOF-MS, liquid chromatography coupled with quadrupole time-of-flight mass spectrometry; RUS, random undersampling; SMOTE, Synthetic Minority Over-sampling Technique; UD, unbalanced data.

To verify the reliability of our findings, the models developed using targeted analysis data were revalidated with an independent subset of samples (see Figure 6). The accuracy values and F1 scores for the RF-based model developed for glioma differed by 0.01–0.03 across the UD subsets and when balancing with SMOTE and RUS. For meningiomas, the F1 scores were, on average, 5.9% higher for models that used the RUS-based class-balancing algorithm. For the model using SMOTE on 10 patients, the F1 score was 5% lower. No significant differences were observed for 20 and 30 patients. When testing the model developed for unbalanced groups, the F1 score varied by -3.6% to 2.8%. Revalidation of SVM-based models for glioma showed an average decrease in F1 score of 9.5% for the RUS-balanced set and around 1% for the SMOTE-balanced set. Unbalanced sets varied in F1 scores from -0.01 to 0.05. For meningioma samples, the F1 score varied by about 3% across all subsets balanced with RUS and SMOTE. With the UD, a decrease in F1 score was observed as the number of real samples used for modeling increased, with percentage changes in F1 score from -6.7% to 0.6%.

**Figure 6.**
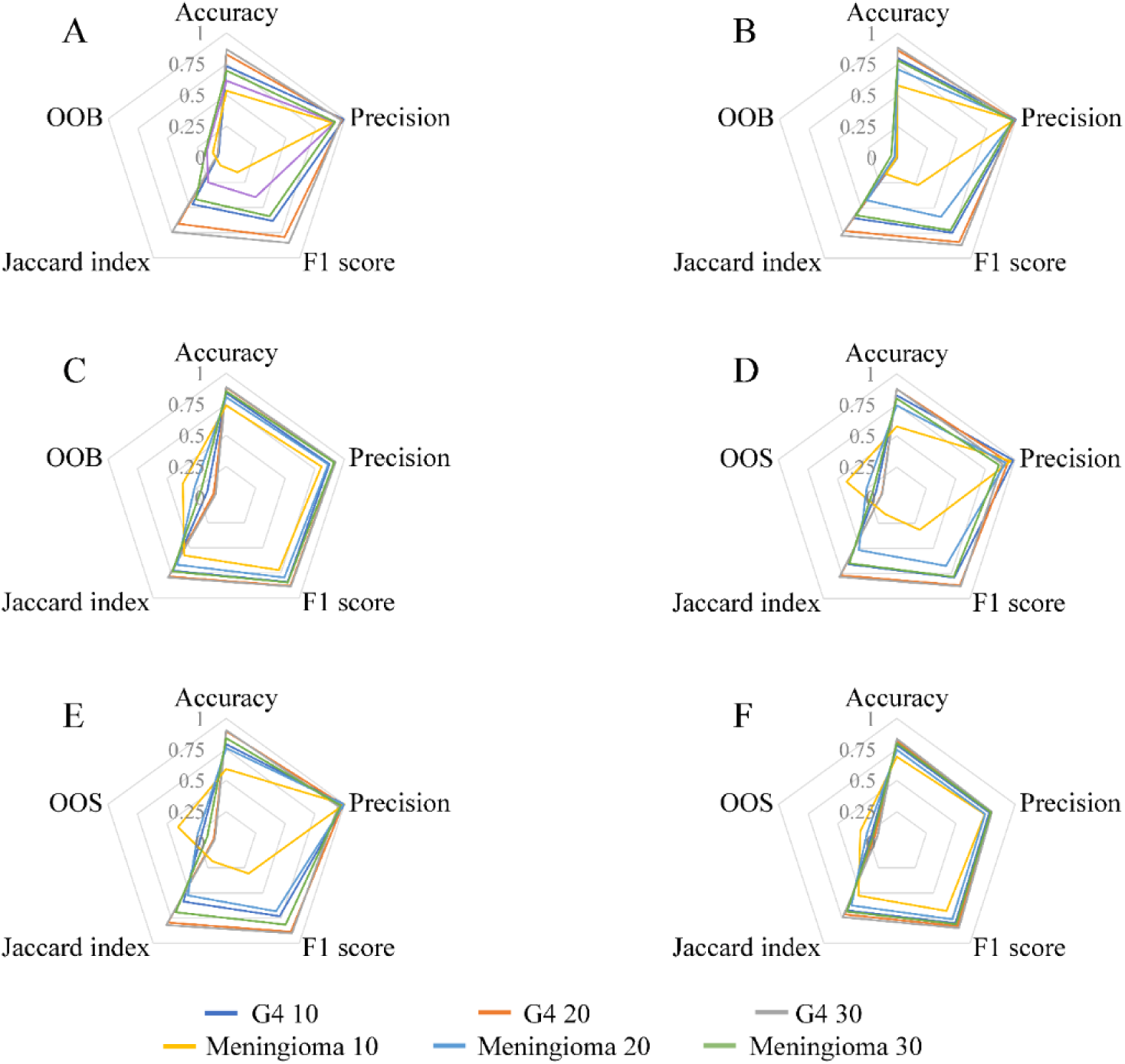
Radar charts showing classification metrics for revalidated models developed using results from targeted analysis: (A) RF model without oversampling; (B) RF model with SMOTE; (C) RF model with undersampling; (D) SVM model without oversampling; (E) SVM model with SMOTE; (F) SVM model with undersampling. OOB, out-of-bag error; OOS, out of sample error.

## 4. DISCUSSION

Metabolomic analysis provides comprehensive insights into metabolic alterations associated with disease progression. The reliability of these analyses depends strongly on sufficiently large and well-balanced study cohorts that minimize biological and environmental variability. However, recruiting comparable numbers of participants across study groups within a single center is often challenging. Although multicenter studies can increase cohort size, they may introduce additional variability due to differences in sampling procedures, analytical platforms, and data-processing workflows. Such heterogeneity may arise during sample collection and preparation, from instrumental performance or inconsistencies in metabolite annotation and nomenclature, ultimately contributing to batch effects and reduced comparability of datasets [30–33]. To address class imbalance, researchers frequently use innovative computational methods to artificially balance study groups. Many existing balancing algorithms, including oversampling and undersampling methods, are well documented in the literature [10, 34–38]. Despite their popularity, the impact of these methods on metabolomic data distortion has not been systematically evaluated. Our study complements this knowledge by assessing the impact of commonly used balancing algorithms on data distribution across three analytical platforms and two cohorts of patients with brain tumors.

The straightforward approach to equalize group sizes is to exclude samples from the majority class. However, manual removal can involve selection bias. The RUS algorithm addresses this issue by randomly removing samples from the majority group. If the class imbalance is substantial (IR above 7.0), applying this method may result in the loss of biologically relevant information [39, 40]. Consequently, oversampling techniques are more frequently employed in omics studies of rare cancers or infrequent disease subtypes [11, 41]. For instance, low-grade gliomas often progress asymptomatically and are frequently detected incidentally during imaging examinations performed for unrelated conditions, which significantly limits the ability to recruit a sufficient study group. Similarly, specialized clinical centers may encounter challenges in recruiting appropriate control groups, further motivating the use of oversampling algorithms [18, 40, 42].

### Univariate statistics

Balancing the number of observations across groups through oversampling increases the cohort size, thereby improving the statistical power of the tests [43]. Our study demonstrated that applying RUS to data with strong class imbalance (n = 10) substantially reduces statistical power, resulting in approximately a threefold decrease in the number of SSM across all analytical platforms used, compared with UD when comparing gliomas and controls (see Figure 2) [35, 40]. In the meningioma dataset (n = 10), no discriminating metabolites were identified after applying this algorithm. Notably, for both types of brain tumors, fewer metabolites distinguishing them from the control group were observed after RUS application than in UD for n = 20 and n = 30. Conversely, employing SMOTE enabled the detection of a considerably greater number of SSM compared to UD. The increase in SSM when comparing brain tumors to controls was closely related to the number of patients used for oversampling and inversely proportional to the initial size of the minority group. This occurs because oversampling a small group generates synthetic samples from neighboring observations, thereby increasing the homogeneity of the minority class [44]. Consequently, this may artificially inflate statistical power and lead to FPs. The number of differentiating features identified after applying SMOTE also depended on the analytical platform used, particularly on the number of metabolites measured. In the LC-QTOF-MS dataset, the data matrix comprised 809 metabolomic features, resulting in the highest number of statistically significant differences, many of which were classified as FPs (see Table 1). However, using SMOTE enabled the detection of nearly the same differences as in the full dataset, regardless of the initial sample size in the minority group.

### Machine learning modeling

Balancing group sizes can help reduce bias during the ML model’s training. In cases of severe class imbalance (IR above 7.0), the model may become biased toward the majority class [45]. Furthermore, balanced data ensures the correct calculation of classification model evaluation metrics, which is crucial for proper model validation [46]. In our study, we demonstrated that the effectiveness of class-balancing techniques depends strongly on the ML algorithm used. For RF, both oversampling and undersampling improved classification performance in the glioma and meningioma groups (see Table 2). The largest increase in the F1 score was observed in the 10-sample groups (IR = 7.10), whereas the improvement was lower in the other studied groups. In the glioma group n = 30, it was about 3.3% with SMOTE and 4.5% with RUS. Interestingly, for models supported by undersampling, no decline in the F1 score relative to UD was observed in the meningioma dataset. Moreover, in both brain tumor groups, RUS yielded higher values than SMOTE during RF modeling. With SVM, using SMOTE resulted in only decreases or fluctuations (average percentage change: -5%) in classification metrics, regardless of the technique or sample type. For RUS, it increased by about 23% on average in the glioma group with n = 10, whereas changes in the n = 20 (IR = 3.55) and n = 30 subsets (IR = 2.37) were minor , amounting to 2.6% and -0.4%, respectively. For meningiomas, a significantly greater improvement in the F1 score was observed in the n = 10 and n = 20 groups, with average increases across all platforms of 155.9% and 17.2%, respectively. In contrast, in the n = 30 group, the average improvement in the F1 score was only about 3%, while the use of RUS contributed to an average 2% decrease in accuracy in the LC-QQQ-MS and GC-MS data. These differences may be attributed to the distinct learning mechanisms of RF and SVM. RF, due to its ensemble structure and reliance on bootstrap aggregation, tends to be relatively robust to distributional shifts introduced by resampling. In contrast, SVM optimizes the decision boundary through margin maximization and is therefore more sensitive to alterations in class geometry induced by synthetic observations [47, 48]. Additionally, removing samples from the majority class using RUS leads to a loss of information about the control population, reducing the representativeness of its distribution [39, 40], which may increase the risk of overfitting. This effect was especially pronounced in meningiomas, where applying RUS increased the F1-score from 0.17 (UD set) to 0.71 in the LC-QTOF-MS dataset.

To confirm our findings, we conducted additional targeted analyses to validate the developed ML methods on an independent test set (see Figure 6). When using RUS to model meningiomas with RF, we observed a 5.8% increase in the F1 score, suggesting a beneficial effect of group balancing. At the same time, a 5% decrease in this metric was noted when using SMOTE for the n = 10 group. This may reflect the introduction of synthetic samples within a limited feature space, which can artificially densify local patterns and reduce model generalizability, leading to overfitting. Interestingly, in SVM modeling, revalidation showed an average decrease in F1 of 9.5% for glioma datasets balanced using the RUS method. This effect likely results from the very small sample sizes after undersampling, which limit the representation of the majority-class distribution and increase model variance. Similar effects have been observed in previous studies, where undersampling reduced model generalization by removing informative samples from the majority class [49]. Conversely, oversampling, particularly in small datasets, tends to exaggerate the importance of features and increases the risk of FPs [10, 50]. Therefore, both undersampling and oversampling should be applied carefully in metabolomic studies, especially in high-dimensional, low-sample-size settings, ensuring adequate cohort size to minimize overfitting and preserve biological variability.

### Distribution and correlation analysis

Additionally, our study examined the impact of both balancing algorithms on data distribution depending on the number of samples and the analytical technique used. The use of RUS preserved the original data distribution for each measured metabolite. In contrast, the use of SMOTE resulted in significant differences for several metabolites (Figure 3). These differences likely reflect changes in local data density introduced by interpolation between neighboring samples rather than true biological variation. For the oversampling of the n = 30 group, fewer metabolites reached significance in the KS test, suggesting that the negative effect of oversampling diminishes with increasing sample size. Interestingly, similar trends were observed for both meningiomas and gliomas, whereas platform-specific effects were evident in GC-MS data. Using SMOTE on the glioma dataset, metabolites became statistically significant in the comparisons, whereas in the case of meningioma comparisons, p-values were higher. This suggests that the influence of oversampling depends primarily on data structure, feature-space geometry, rather than on tumor type alone [51].

We confirmed our observations by conducting SW tests (Supplementary Tables S1 and S2) and by visualizing the results with violin plots. (Figure 4). The selected metabolites had distinct data distributions. Leucine, measured by GC-MS, exhibited an approximately normal distribution in the full dataset, whereas the glioma dataset (n = 10) produced a non-normal distribution after SMOTE application. As the sample size increased, the p-value of the SW test gradually converged toward the value observed in the original dataset, suggesting that the distributional distortion caused by the SMOTE algorithm was more pronounced under severe class imbalance. After applying SMOTE, the leucine distribution shifted toward non-normality across all meningioma groups studied. Conversely, for proline (LC-QQQ-MS) and the metabolic feature with a neutral mass of 607.3351 (LC-QTOF-MS), no change in data distribution was observed with oversampling. This is related to the observation that most data from actual measurements exhibit a skewed distribution [52]. SMOTE generates new samples through linear interpolation between neighboring observations, which does not guarantee preservation of the original multivariate distribution. In high-dimensional metabolomic data, this may lead to local covariance shrinkage, smoothing of natural variability, and the generation of synthetic observations that do not fully capture underlying biological heterogeneity. Furthermore, data noise and outliers can generate unrepresentative synthetic samples, with extreme values distorting the data structure [51]. Consequently, the generated data may amplify the influence of outliers, thereby negatively affecting the reliability of statistical analyses and modeling. Conversely, RUS does not introduce artificial data and therefore preserves the original feature relationships among retained observations while randomly removing samples from the majority class, which may also affect the data distribution [22].

Comparing the heatmaps generated for metabolites measured using both LC-QQQ-MS and GC-MS, we observed disruptions caused by the SMOTE algorithm (see Figure 5). Oversampling almost completely altered the structure of glioma and meningioma data (n = 10), eliminating correlations between platforms. Interestingly, only when n = 30 was the correlation strength comparable to that observed in the full dataset, though not all were statistically significant. This effect was related to changes in covariance structure introduced by SMOTE, as well as variability inherent in generating synthetic samples across iterations [10]. Sequentially generated synthetic samples for the LC-QQQ-MS and GC-MS datasets differ in the interpolation coefficient (a random number between 0 and 1 used in the SMOTE formula), introducing minor random variation. As the number of synthetic samples increases, correlation coefficients between platforms may decrease. Loss of inter-platform correlation can hinder integration of metabolomic and clinical data, potentially leading to biased or erroneous interpretations [53].

### Practical guidelines

The conditions required for proper data balancing have been outlined based on the literature [44, 50, 54–60] and our knowledge. After conducting the experiment in accordance with analytical standards, the data quality should be carefully assessed, followed by the exclusion of all metabolites that do not meet the specified criteria [54]. This process reduces noise and decreases the number of features that could distort the results. Subsequently, data preprocessing steps such as normalization, missing value imputation, and, if necessary, batch effect correction should be applied to ensure consistency and minimize technical variability [55, 56].

Before conducting statistical analysis, the choice of balancing algorithm must be carefully considered. Simple duplication of minority class observations should be avoided, as it can lead to overfitting. Oversampling is recommended for smaller datasets where each sample is valuable, and the disproportionate group sizes could negatively affect model generalization. Conversely, undersampling is more appropriate for large datasets, where removing samples from the majority class is unlikely to significantly affect model performance. In cases of high class imbalance (IR above 7.0), combined approaches (e.g. SMOTE+RUS) should be considered, as reliance on a single method may lead to overfitting with SMOTE or reduced classification performance with RUS [44, 57]. Maintaining a minimum number of samples is also essential to ensure reliable statistical inference and robust ML model training, especially in high-dimensional metabolomic datasets.

When performing univariate statistical testing, multiple iterations with different sample subsets should be conducted, allowing p-values to be averaged and thus reducing the likelihood of FPs. Appropriate tests, such as the SW, KW, or t-test, should be employed to determine whether synthetic samples differ significantly from the original dataset [58].

Prior to training the ML models, the number of variables should be reduced by removing irrelevant or redundant metabolites [59]. Metabolite selection methods, such as LASSO or mRMR (minimum-Redundancy-Maximum-Relevance), can minimize the inclusion of irrelevant variables into the synthetic sample generation process, thereby reducing potential bias [59, 60]. A crucial consideration in oversampling-assisted modeling is ensuring that synthetic samples are confined to the training set to avoid data leakage, which can artificially inflate classification metrics [50]. Evaluation on real samples provides a more reliable assessment of model performance. Furthermore, robust metrics such as F1 score, AUC–ROC, and the Brier score should be used, as they are less sensitive to unequal class distribution and more informative for imbalanced, high-dimensional datasets [50].

## 5. CONCLUSIONS

In our study, we investigated the effects of oversampling and undersampling on plasma metabolomic data from patients with glioma and meningioma, and from controls. All experiments were performed using three complementary techniques, enabling us to thoroughly evaluate the performance of balancing algorithms across different data types. While algorithms that address class imbalance can be applied in metabolomic studies, careful consideration is necessary when using them. They may increase FP, cause model overfitting, and reduce correlation. However, in certain cases, using SMOTE may be beneficial, as we demonstrated that even a small number of samples can yield the same SSM as the full dataset. This approach can be particularly useful in statistical analyses aimed at selecting metabolic features for annotation or metabolites for validation. Models developed using ML algorithms should be validated with an independent experiment to avoid overfitting. The guidelines we have provided can reduce the adverse effects of data balancing. However, it is important to carefully consider when these algorithms are necessary and to critically evaluate the obtained results.

