## Supplementary Materials for "Application of class–balancing algorithms to diverse plasma metabolomics datasets using brain tumor as an example"

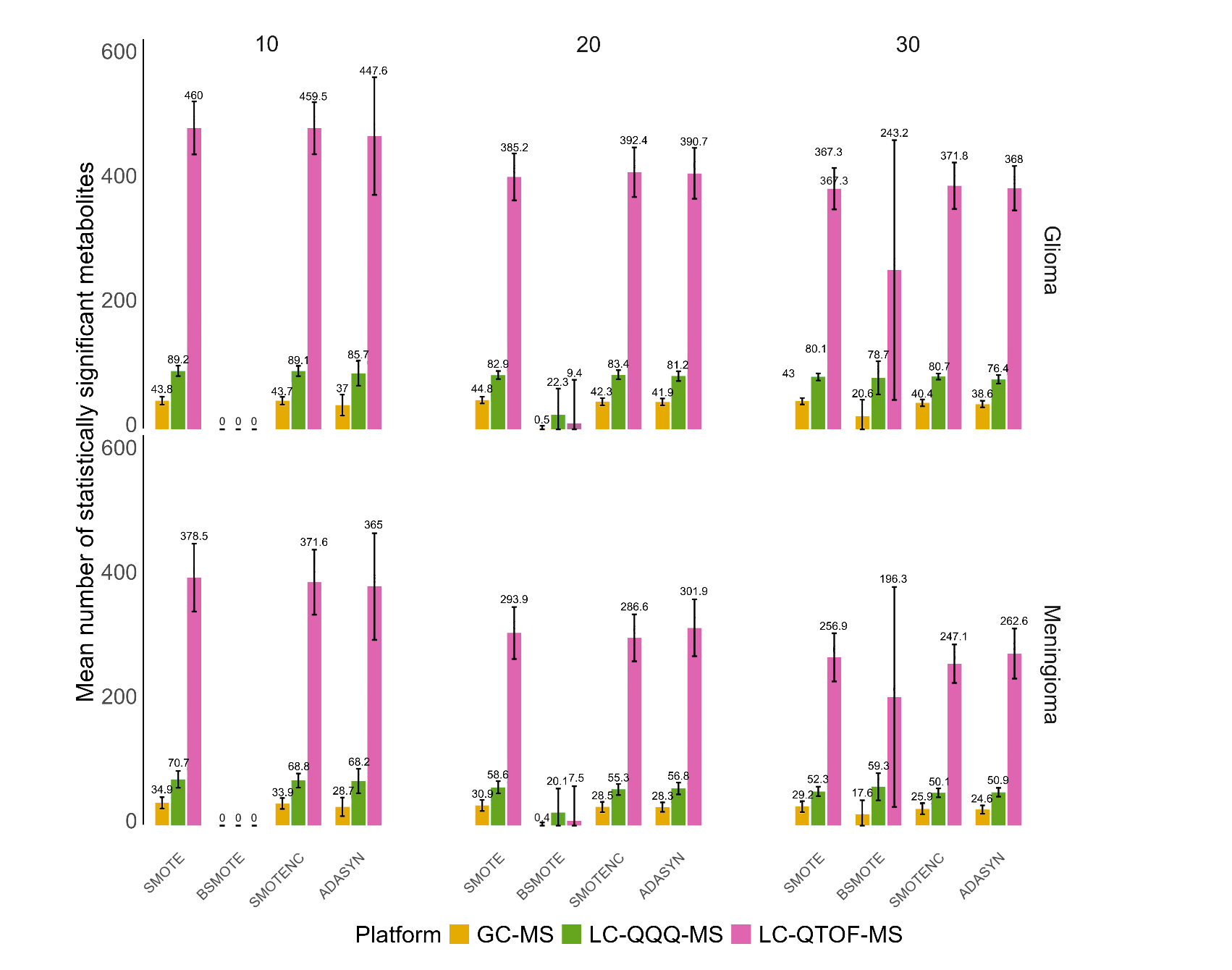

Supplementary Figure S1. The average number of SSM for GC-MS, LC-QQQ-MS, and
LC-QTOF-MS, across 100 iterations, considering subgroup division and the application
of oversampling algorithms. ADASYN, Adaptive Synthetic Sampling Approach; BSMOTE, borderline-SMOTE; GC-MS, gas chromatography coupled with mass spectrometry;
LC-QQQ-MS, liquid chromatography coupled with tandem mass spectrometry; LC-TOF-MS, liquid chromatography coupled with quadrupole time-of-flight mass spectrometry; SMOTE, Synthetic Minority Over-sampling Technique; SMOTENC, SMOTE for Numeric and Categorical data; UD, unbalanced data.

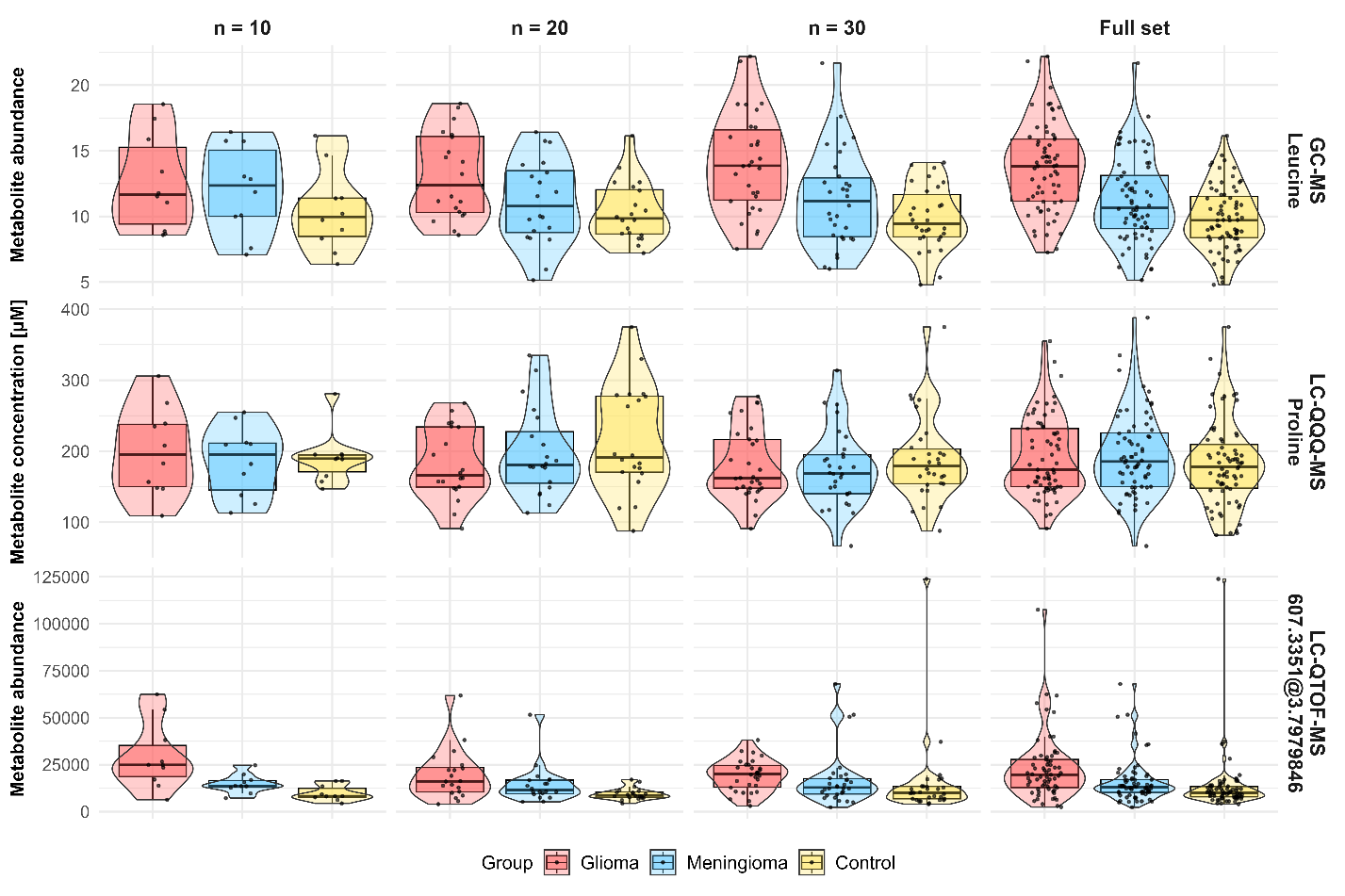

Supplementary Figure S2. Violin plots representing the abundance of leucine measured
by GC-MS, the concentration of proline measured by LC-QQQ-MS, and the abundance of the metabolic feature with a neutral mass of 607.3351 Da measured by LC-QTOF-MS after balancing the groups using RUS. GC-MS, gas chromatography coupled with mass spectrometry;
LC-QQQ-MS, liquid chromatography coupled with tandem mass spectrometry; LC-TOF-MS, liquid chromatography coupled with quadrupole time-of-flight mass spectrometry.
